# Monitoring intracellular antibiotic concentrations in real-time using allosteric biosensors

**DOI:** 10.64898/2026.02.05.704027

**Authors:** Daria Fleckenstein, Andreas Kaczmarczyk, Niklas Breitenbach-Netter, Gili Rosenberg, Roman Peter Jakob, Isabel Sorg, Amanzhol Kurmashev, Carlos Flores, Eva Jiménez-Siebert, Elinor Morris, Steffi Klimke, Sarah Tschudin-Sutter, Andreas Hierlemann, Timm Maier, Christoph Dehio, Urs Jenal, Knut Drescher

**Author notes:** Equal contribution (Daria Fleckenstein, Andreas Kaczmarczyk).

## Abstract

Antibiotic treatment can fail due to insufficient drug availability at the site of infection or limited accumulation within bacterial pathogens. However, it is poorly understood how antibiotics penetrate infected tissues and complex bacterial aggregates, limiting insights into the mechanisms of treatment failure. Here, we present genetically-encoded allosteric biosensors for two antibiotic classes, trimethoprim and tetracycline, which enable real-time monitoring of antibiotic concentrations inside bacterial cells. The biosensors consist of circularly permuted EGFP linked to the sensory domains DHFR or TetR. To extend this approach to low oxygen environments, we engineered an oxygen-independent trimethoprim biosensor by fusing DHFR to a circularly permuted version of the fluorogenic protein FAST. Using these biosensors, we monitored the antibiotic exposure dynamics of intracellular *Salmonella enterica* during macrophage infection at the single-cell level, and antibiotic penetration into anaerobic regions of *Vibrio cholerae* biofilms, as well as antibiotic availability in microoxic conditions in a human bladder tissue model infected with uropathogenic *Escherichia coli*. These fluorescent biosensors have the potential to be broadly applied for determining antibiotic distributions at infection sites with high spatial and temporal resolution.

## Introduction

Since their introduction into medical practice, antibiotics have revolutionized healthcare and saved countless human lives. Unfortunately, antibiotic treatment of infections often fails, with critical consequences for patients. Understanding the mechanisms underlying treatment failure is of major importance for designing new drugs and for adapting therapeutic strategies with existing antibiotics. Treatment can fail if a bacterial strain is resistant to an antibiotic and continues to grow in a particular concentration range of the antibiotic. Treatment can also fail when bacterial strains tolerate specific antibiotic doses, or when drug concentrations at the infection site remain too low to kill or inactivate bacteria. Key determinants of treatment success or failure are the intracellular antibiotic concentration at the site of infection, and the duration for which an elevated intracellular antibiotic concentration is achieved.

Current methods for assessing intracellular antibiotic concentration dynamics have significant limitations. Some methods lack spatial resolution (*e*.*g*., disc diffusion assays^1–3^, radiolabeling^4^, mass spectrometry^5^), or have spatial resolution but low temporal resolution and only indirectly reflect the antibiotic concentration (GFP-based transcriptional reporters^6,7^ or measuring single-cell morphological changes^8,9^), are technically highly demanding (e.g., Raman spectroscopy^10^, attenuated total reflectance Fourier transform infrared spectroscopy^11^), or involve labeling of the antibiotic with a fluorescent dye which alters the antibiotic function and transport properties^12–15^.

To overcome these limitations, we have developed genetically-encoded allosteric biosensors capable of real-time detection of unmodified antibiotics inside live bacterial cells. We developed both O_2_-dependent and O_2_-independent biosensor variants, which enable measurements of intracellular antibiotic concentrations at high spatial and temporal resolution using fluorescence microscopy.

## Results and discussion

### Design and characterization of antibiotic biosensors

The genetically encoded allosteric biosensors comprise two parts: a sensory unit that binds the antibiotic of interest, and a fluorescence unit that undergoes a change in fluorescence depending on the analyte binding (Fig. 1a,b). As a fluorescence unit, we chose circularly permuted EGFP (cpEGFP^16^). As sensory unit candidates, we considered all proteins with significant structural changes between the antibiotic-bound and unbound states based on structures in Protein Data Bank (PDB). For sensory unit candidates with enzymatic activity directed to cleave or modify the antibiotic, we introduced inactivating amino acid substitutions in their catalytic motifs.

**Fig. 1.**
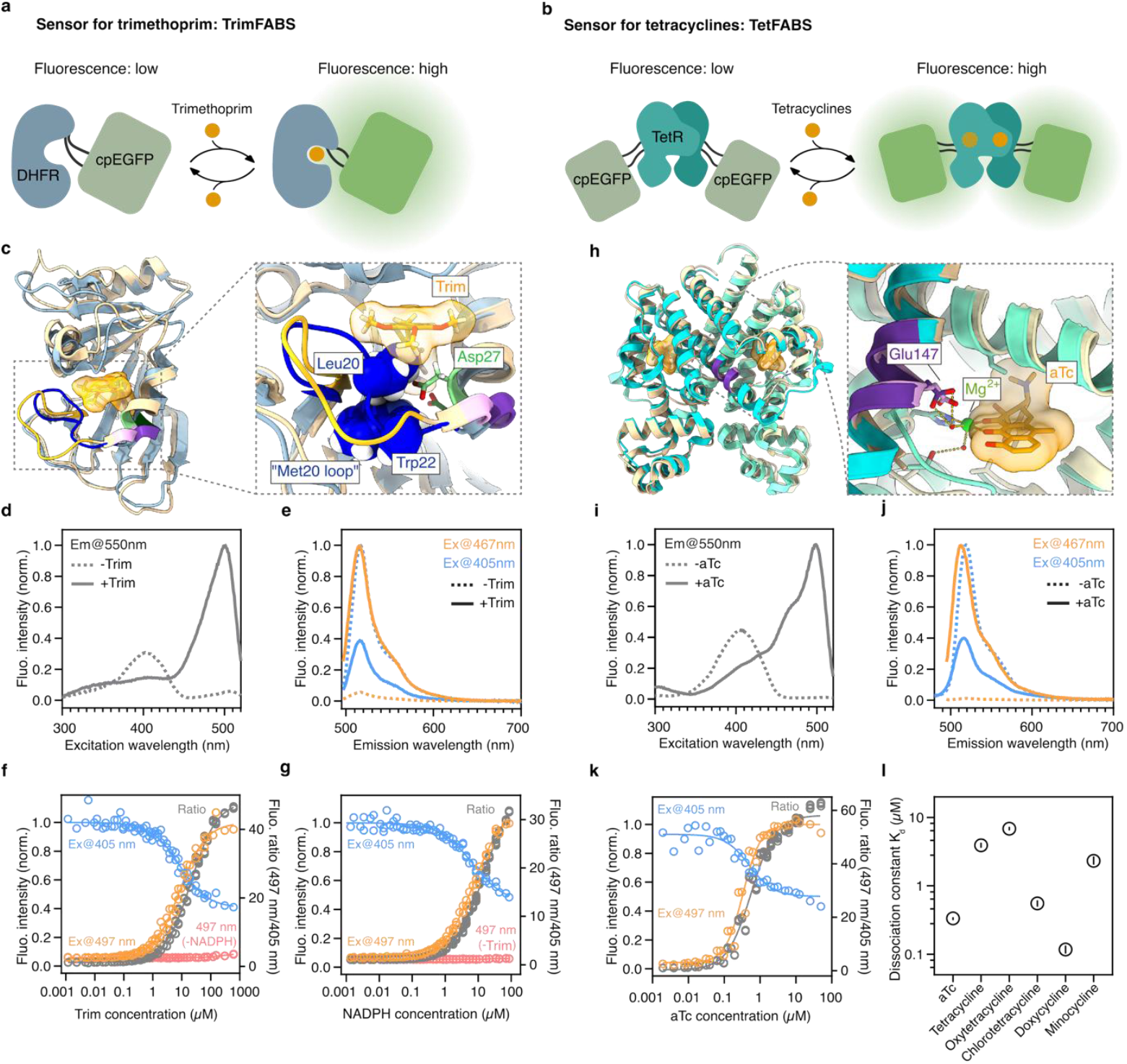
Design and *in vitro* characterization of antibiotic biosensors. **a-b**, Schematic representation of the genetically encoded allosteric biosensors for (a) trimethoprim and (b) tetracyclines. **c**, Overlay of DHFR in the trimethoprim (Trim)-bound (blue-gray) state and unbound state (beige), in cartoon presentation (left) with a close-up of the Trim binding site (right). The “Met20 loop” in the apo- and holo-form is highlighted in bright yellow and dark blue, respectively; the active site Asp27 is highlighted in shades of green; the insertion site of cpEGFP in shades of violet. In the close-up view, hydrophobic residues Leu20 (corresponding to the namesake Met20 in *E. coli*) and Trp22 in the “Met20 loop” that flip into the binding pocket upon Trim-binding are displayed as spheres in the holo-structure. **d**, Excitation spectra of trimethoprim sensor TrimFABS with and without Trim in the presence of 250 μM NADPH. **e**, Emission spectra of TrimFABS with and without Trim and excitation at 405 nm (blue) or 467 nm (orange) in the presence of 250 μM NADPH. **f**, Trim dose-response curves of TrimFABS in the presence of 250 μM NADPH, showing emission at 550 nm with excitation at 405 nm or 497 nm (left y-axis) and the combined ratiometric readout (gray points, ratio of 550 nm emission signal with 497 nm and 405 nm; right *y*-axis). A Trim dose-response curve of TrimFABS without NADPH, excited at 497 nm, is shown for comparison (red). Data points are pooled from n=6 independent dilution series with six repeated measurements for each concentration at steady state. **g**, NADPH dose-response curves of TrimFABS in the presence of 1.25 mM Trim, showing emission at 550 nm with excitation at 405 nm and 497 nm (left y-axis) and the combined ratiometric readout (ratio of 550 nm emission signal with 497 nm and 405 nm; right *y*-axis). A NADPH dose-response curve of TrimFABS without Trim at 497 nm excitation is shown for comparison (red). Data points represent pooled from n=6 independent dilution series with six repeated measurements for each concentration at steady state. **h**, Overlay of TetR in the anydrotetracycline (aTc)-bound (turquoise) and unbound state (beige), in cartoon presentation (left) and a close-up of the binding site (right). The region (residues 145-149) where cpEGFP* is inserted is highlighted in light and dark violet in the aTc-unbound and -bound forms, respectively. The crucial Glu147 (shown as stick) and magnesium ion (green sphere) stabilizing tetracycline binding via water molecules (small red spheres) are indicated. **i**, Excitation spectra of tetracycline sensor TetFABS with and without aTc. **j**, Emission spectra of TetFABS with and without aTc and excitation at 405 nm (blue lines) or 467 nm (orange lines). **k**, aTc dose-response curves of TrimFABS emission at 550 nm with excitation at 405 nm or 497 nm (left y-axis) and the combined ratiometric readout combined ratiometric readout (gray points, ratio of 550 nm emission signal with 497 nm and 405 nm; right *y*-axis). Data points represent pooled data from n=3 independent dilution series with six repeated measurements for each concentration at steady state. **l**, Affinity (K_d_) of different antibiotics of the class of tetracyclines to TrimFABS. Shown are mean (circles) and 95% confidence intervals (lines) obtained from fits of dose-response curves, from n=3-6 independent replicas for each condition (see Supplementary Fig. 1).

To construct biosensor candidates from the combination of the sensory and fluorescent units in an unbiased manner, we implemented a transposon-based random domain-insertion library method^16^ in which cpEGFP was inserted at random locations of the sensory unit sequence. Using iterative fluorescence-activated cell sorting (FACS) with and without the antibiotic of interest, we enriched for functional biosensors. As a result, we obtained functional biosensors for trimethoprim (based on NADPH-dependent dihydrofolate reductase, DHFR) and for tetracyclines (based on the tetracycline repressor, TetR). These first-generation biosensors were then optimized by randomizing linker regions between the sensory and fluorescence units, followed by iterative FACS to select better-performing biosensors, which resulted in the final biosensors reported and characterized in this study (Fig. 1a,b).

The allosteric biosensor for trimethoprim (TrimFABS – <u>trim</u>ethoprim fluorescent <u>a</u>nti<u>b</u>iotic <u>s</u>ensor) has cpEGFP inserted in direct vicinity of the catalytic Asp27 and following the flexible Met20 loop near the substrate binding pocket of DHFR^17^. This region of DHFR undergoes a conformational change upon trimethoprim binding and is therefore suitable for inducing a fluorescent cpEGFP conformation (Fig. 1c). Purified TrimFABS in the presence of the cofactor NADPH showed two excitation maxima with and without trimethoprim at 499 nm and 402 nm, respectively, with a shared single emission peak at 516 nm (Fig. 1d,e). The inverse behavior of the two excitation peaks upon ligand binding enable a ratiometric readout. Trimethoprim dose-response curves revealed a dose-dependent, monotonic behavior for both major excitation wavelengths, with a fold change of ∼17.1 and a fitted K_d_ of 8.7 μM (95% CI: 8.3-9.2 μM; Hill slope: 0.97) at 497 nm and a fold change of ∼2.5 and a fitted K_d_ of 8.4 μM (95% CI: 7.5-9.4 μM; Hill slope: -0.88) at 405 nm, resulting in a combined ratiometric fold change of ∼42 and a fitted K_d_ of 20.9 μM (95% CI: 20.0-22.0 μM; Hill slope: 0.96) (Fig. 1f).

The response of TrimFABS to trimethoprim strictly depends on NADPH, which is known to be required for binding of the natural DHFR substrate dihydrofolate^17^ (Fig. 1f). Likewise, TrimFABS responded to NADPH in a dose-dependent manner in the presence of trimethoprim when excited at 497 nm or 405 nm (Fig 1g), with fold changes of ∼16 or ∼2.3, respectively, and K_d_ of 7.4 μM (95% CI: 7.2-7.7 μM; Hill slope: 0.98) or 6.7 μM (95% CI: 5.5-8.6 μM; Hill slope: -0.86), respectively. The combined ratiometric readout for NADPH binding yields a predicted maximum fold change of ∼36 and a fitted K_d_ of 15.2 μM (95% CI: 14.7-15.8 μM; Hill slope: 0.99) (Fig 1g). Given that intracellular NADPH concentrations are typically in the range of 100-200 μM^18–20^ and thus well above the K_d_ for NADPH, these results suggest that TrimFABS is saturated with NADPH *in vivo*, therefore accurately and exclusively reporting on trimethoprim levels.

In contrast to TrimFABS, the biosensor for tetracyclines (TetFABS – <u>tet</u>racyclines fluorescent <u>a</u>nti<u>b</u>iotics <u>s</u>ensor) has cpEGFP inserted in a rigid α-helical region that is not predicted to undergo major conformational changes (Fig. 1h), but is adjacent to Glu147, which – via a water molecule – coordinates a Mg^2+^ ion that stabilizes the binding of tetracyclines^21^. The side chain of Glu147 was shown to rotate upon antibiotic binding^21^. This minor conformational change seems to translate into major alterations in fluorescence intensity of the cpEGFP moiety. TetFABS has two excitation maxima at 498 nm and 404 nm with corresponding emission peaks at 512 nm and 519 nm, respectively, which respond inversely to the addition of anhydrotetracycline (Fig. 1i,j). Dose-response curves recorded with anhydrotetracycline indicate a dose-dependent, monotonic behavior for the two excitation wavelengths, with a fold change of ∼23 and a fitted K_d_ of 335 nM (95% CI: 314-357 nM; Hill slope: 1.8) at 497 nm and a fold change of ∼1.9 and a fitted K_d_ of 502 nM (95% CI: 433-588 nM; Hill slope: -1.3) at 405 nm, resulting in a combined ratiometric fold change of ∼43 and a fitted K_d_ of 571 nM (95% CI: 532-615 nM; Hill slope: 1.3) (Fig. 1k). Other antibiotics from the class of tetracyclines show similar dose-dependent monotonic behaviors with varying affinities (Fig 1l, Supplementary Fig. 1), suggesting that TetFABS is a versatile tool for the detection of different clinically relevant tetracyclines.

Using the dissociation-by-dilution method we determined k_off_ rate constants of 3.32 min^-1^ (95% CI: 3.01-3.66) and 4.65 min^-1^ (95% CI: 4.01-5.38) for TrimFABS and TetFABS, respectively. This corresponds to binding half-lives of 0.21 min (95% CI: 0.19-0.23 min) (TrimFABS) and 0.15 min (95% CI: 0.13-0.17 min) (TetFABS) (Supplementary Fig. 2a,c). Based on these data, we calculate association rate constants (k_on_ = k_off_ K_d_^-1^) of 4.5 × 10^5^ M^-1^ min^-1^ for TrimFABS and 4.0 × 10^7^ M^-1^ min^-1^ for TetFABS. To complement these data and provide a reference for *in vivo* experiments, we also determined the time needed to obtain 50% of the maximal signal after addition of ligand in the pseudo-first-order regime, *i*.*e*., saturation halftime (t_1/2max_). This yielded t_1/2max_ values of 2.3 min for TrimFABS and 2.7 min for TetFABS with doxycycline (Supplementary Fig. 2 b,d), which is at least an order of magnitude faster than the bacterial doubling time. These data show that both biosensors respond with similar rapid kinetics to their corresponding ligands and are thus suitable to study antibiotics dynamics *in vivo* on a physiologically relevant timescale.

### Real-time measurements of antibiotic penetration into intracellular bacteria

Different antibiotics have variable potency to breach through membranes of human host cells, resulting in different antibiotic exposure of intracellular bacteria^22–26^. To reveal the local concentration dynamics of antibiotics inside intracellular bacteria, we infected macrophages with *Salmonella enterica* producing TetFABS or TrimFABS, in addition to a constitutively produced mScarlet-I that was used for image segmentation and fluorescence intensity normalization (Fig. 2a). Using confocal microscopy, we observed that both doxycycline and trimethoprim were able to reach all *S. enterica* cells inside the macrophages (Fig. 2b). Notably, doxycycline takes significantly longer than trimethoprim to enter the intracellular bacteria (Fig. 2c), suggesting that the macrophage lipid membrane is a significant barrier for doxycycline, but not for trimethoprim.

**Fig. 2.**
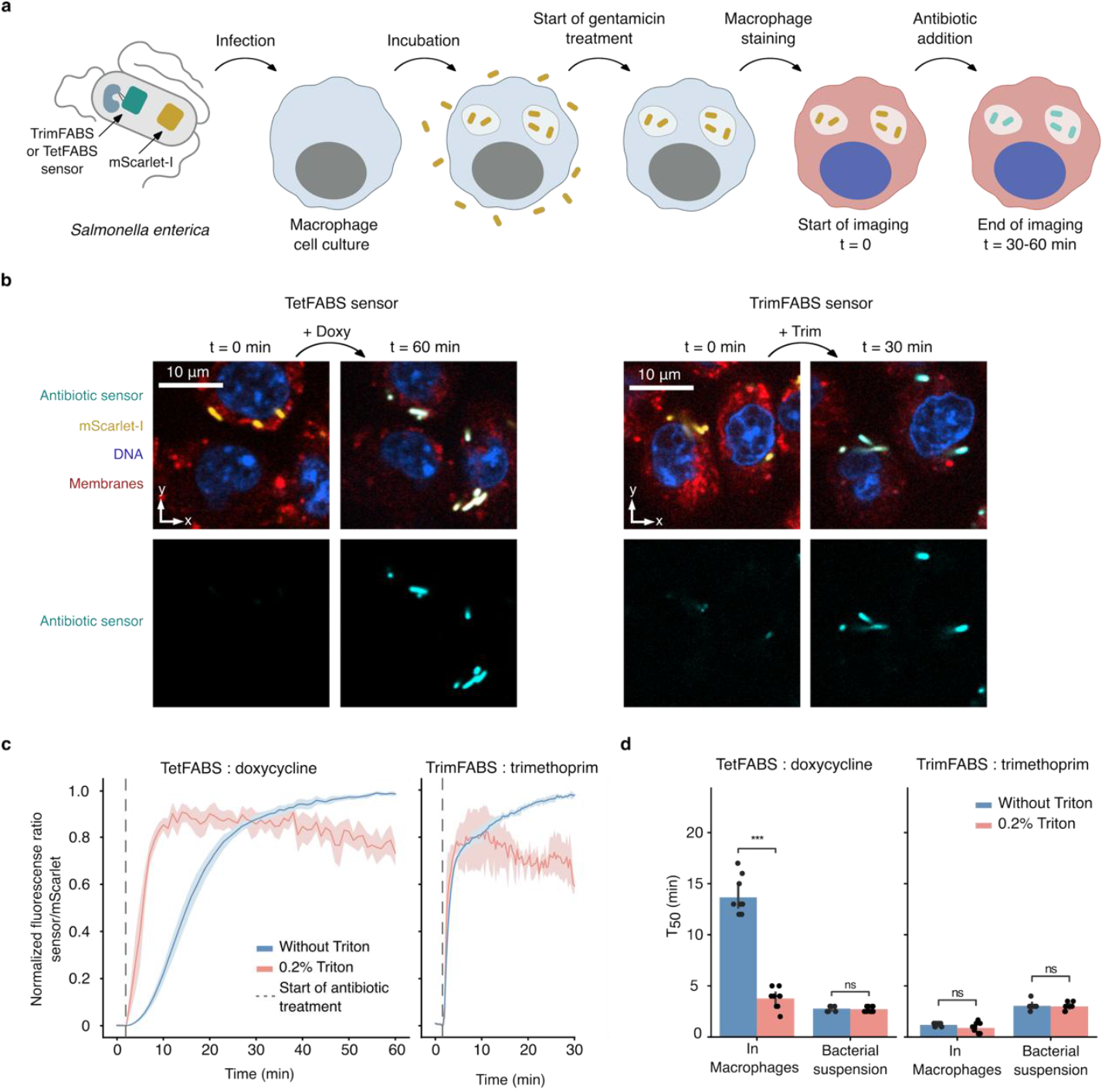
TetFABS and TrimFABS reveal doxycycline and trimethoprim transport dynamics to *S. enterica* inside macrophages. **a**, Schematic of the experiment: RAW macrophages were infected with *S. enterica* harboring TrimFABS or TetFABS with mScarlet. After 30 min incubation, the culture was treated with 50 μg/mL gentamicin to kill non-internalized bacteria, followed by staining macrophages with Hoechst 3342 (nuclei, blue) and CellTracker Deep Red (membranes, red) and time-lapse imaging after the addition of doxycycline (Doxy) or trimethoprim (Trim). **b**, Images of the same field of view before and after Doxy or Trim treatment in four channels: GFP biosensor (cyan), mScarlet (yellow), DNA (blue), membranes (red). **c**, Normalized fluorescence ratio (biosensor/mScarlet) increase over time for of TetFABS after 100 μM Doxy addition and TrimFABS after 100 μM addition in presence (red) or absence (blue) of 0.2% triton. Data points represent pooled data from three positions recorded per well in total of n=3 independent replicas per condition. **d**, Half-saturation time (T_50_) for TetFABS and TrimFABS without triton (blue) or after 0.2% triton treatment in macrophages recorded via microscopy or in control conditions without macrophages measured in bacterial suspension in a plate reader. Data points represent data from three positions recorded per well for n=3 independent replicas. Statistical significances were calculated between triton-treated and untreated conditions using a two-way Mann-Whitney-Wilcoxon test; ns = not significant; * = p < 0.05; ** = p < 0.01; *** = *p* < 0.001; **** = p < 0.0001.

To investigate if antibiotic penetration is indeed limited by the lipid membrane of macrophages, we supplied 0.2% Triton X-100 together with antibiotics, causing macrophages to burst within 20-60 sec, while leaving bacteria intact^27^. Triton-induced membrane damage led to a significant reduction of the time required for doxycycline to enter *Salmonella* (T_50_ dropped from 13.7 ± 1.9 min to 3.8 ± 1.0 min, *p* = 0.00036), but did not alter trimethoprim penetration (Fig. 2d). Triton treatment had no effect on the penetration of the two antibiotics into bacterial cells in pure bacterial cultures, confirming that the macrophage membrane indeed presents a transport barrier specifically for doxycycline (Fig. 2d).

### Development and characterization of an anoxic trimethoprim biosensor

Biofilms are highly abundant communities of bacterial cells embedded in a self-produced extracellular matrix. Compared to planktonic bacteria, biofilms display a strongly increased antibiotic tolerance, which has been attributed to limited penetration of antibiotics into biofilms due to the matrix^1,2,11,28^. Because oxygen is rapidly consumed by cells at the biofilm boundary, the center of large biofilms is generally anoxic, preventing the oxygen-dependent maturation of sensors that are based on GFP-like proteins. To monitor drug penetration into cells deep within biofilms, we engineered a variant of the trimethoprim biosensor in which we replaced cpEGFP with the oxygen-independent circularly permuted fluorescence-activating and absorption-shifting tag (cpFAST) followed by randomizing linkers and using FACS to select the best-performing biosensor, which we named TrimFABS_ANOX_(Fig. 3a). FAST proteins bind externally supplied small molecules called fluorogens (*e*.*g*. HMBR^29^, HBR-3,5DM^30^, HBR-3,5DOM^30^), which are nearly non-fluorescent on their own, but become highly fluorescent in the chemical environment inside FAST proteins upon binding^29^.

**Fig. 3.**
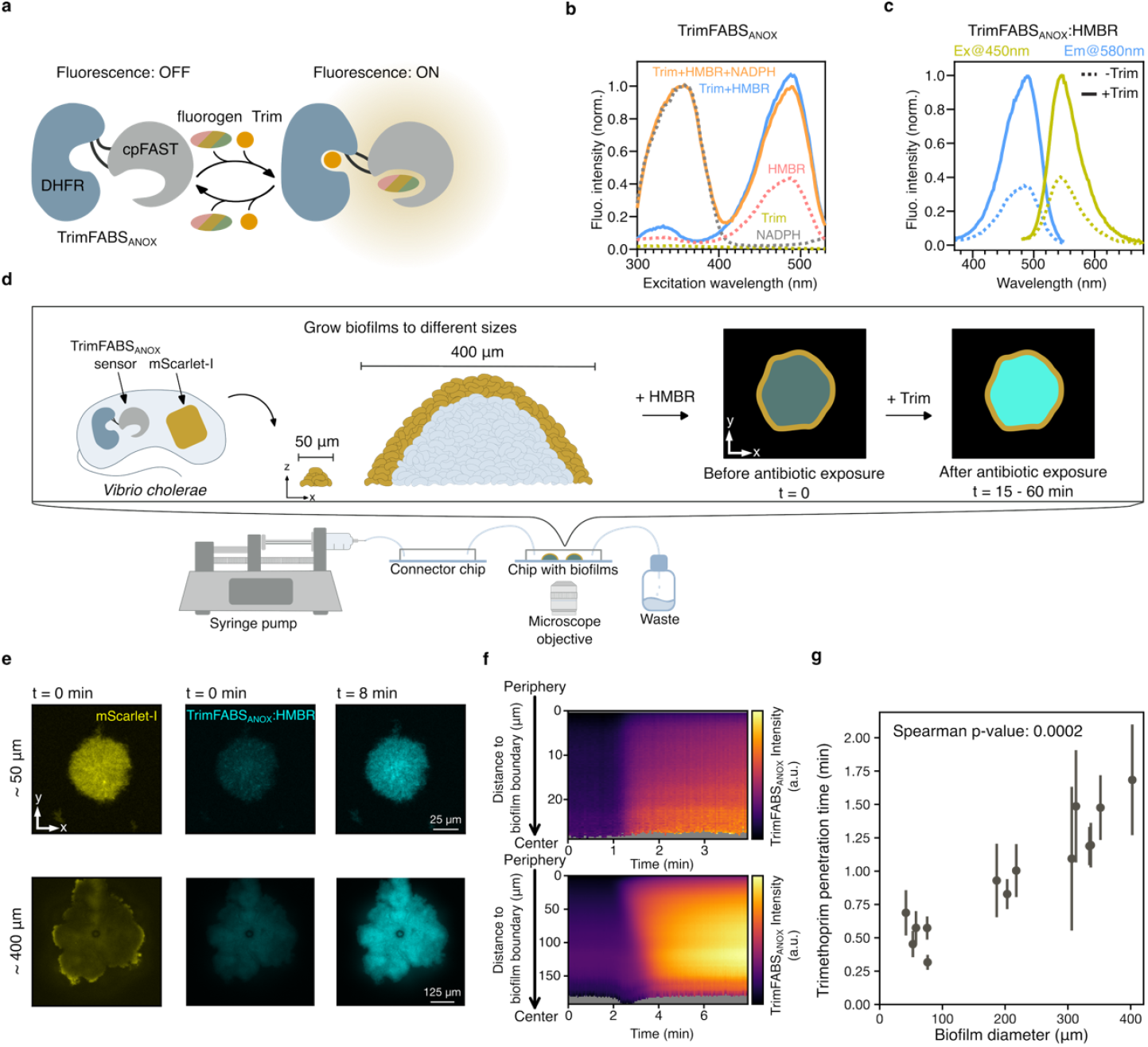
Oxygen-independent biosensor TrimFABS_ANOX_ enables tracking of trimethoprim penetration into bacterial biofilms. **a**, Schematic of TrimFABS_ANOX_: sensory unit DHFR (blue) coupled through linkers (black) with cpFAST (grey) at the same position where cpEGFP is located in TrimFABS. TrimFABS_ANOX_ increases fluorescence upon simultaneous binding of trimethoprim (red) and a fluorogen of choice (multicolored). **b**, Excitation spectra of TrimFABS_ANOX_ with different combinations of Trim, NADPH, and the FAST fluorogen HMBR. **c**, Excitation and emission spectra of TrimFABS_ANOX_ with HMBR as the fluorogen, with or without Trim. **d**, Schematic representation of the experiment: *V. cholerae* strain harboring TrimFABS_ANOX_ and mScarlet on a plasmid was grown in microfluidic device with constant supply of fresh medium until reaching a diameter of 50-500 μm. Then, media was exchanged to supply 5 μM HMBR until sensor saturation, followed by another media exchange to add 25 μM trimethoprim and time-lapse imaging. **e**, Images of biofilms of two different diameters (top row: 50 μm, bottom row: 400 μm) before antibiotic exposure at t = 0 min: mScarlet (yellow, left column) and TrimFABS_ANOX_ (cyan, central column). Images also show TrimFABS_ANOX_ fluorescence at t = 8 min after antibiotic exposure (right column). **f**, Spatiotemporal kymograph of TrimFABS_ANOX_ fluorescence intensity in the ∼50 μm diameter biofilm (top), and ∼400 μm biofilm (bottom). **g**, Trimethoprim penetration time represents the time that is necessary for trimethoprim to reach half saturation in the center of the biofilm after antibiotic addition assuming that sensor response is immediate. Each data point represents mean trimethoprim penetration time across one biofilm. Error bars indicate the standart deviation.

As for other FAST proteins, TrimFABS_ANOX_ excitation is dependent on the presence of a fluorogen, and showed no excitability when incubated with NADPH or trimethoprim alone (Fig. 3b). Fluorescence increased upon trimethoprim binding for several commonly used fluorogens, with fold changes in the excitation peak of 2.9, 1.9, and 4.0 for HMBR^29^, HBR-3,5DM^30^, and HBR-3,5DOM^30^, respectively (Fig. 3c, Supplementary Fig. 3a,b). The spectral properties of the fluorogens were retained in the context of TrimFABS_ANOX_, except that the maximum excitation (λ_ex,max_) and emission (λ_em,max_) were generally red-shifted by 5-10 nm compared to non-permutated FAST proteins (HMBR: λ_ex,max_=490 nm, λ_em,max_=546 nm; HBR-3,5DM: λ_ex,max_=499 nm, λ_em,max_=564 nm; HBR-3,5DOM: λ_ex,max_=530 nm, λ_em,max_=602 nm). Surprisingly, the response of TrimFABS_ANOX_ to trimethoprim was largely independent of NADPH (Fig. 3b), with NADPH only causing a small shift of the fluorescence fold change from 3.0 to 3.5 and the fitted K_d_ for trimethoprim from 6.9 μM (95% CI: 6.1-7.9 μM; Hill slope: 0.7) to 850 nM (95% CI: 710-1000 nM; Hill slope: 0.6) (Supplementary Fig. 3c). These results suggested that TrimFABS_ANOX_ – alone or in complex with a fluorogen – provides new contacts that allow binding of trimethoprim without NADPH and with similar affinities as shown for TrimFABS. Because NADPH still enhances affinity, we presume that these additional interactions do not fully overlap with, mimic, or eliminate NADPH binding. Dose-response curves of the fluorogen HMBR with TrimFABS_ANOX_, with and without trimethoprim, revealed that the antibiotic shifts the apparent K_d_ for HMBR from 2.0 μM (95% CI: 1.8-2.1 μM) to 310 nM (95% CI: 290-330 nM), suggesting that trimethoprim and fluorogen binding are interdependent (Supplementary Fig. 3d). Kinetic characterization of TrimFABS_ANOX_ revealed that the saturation halftime is very rapid (<10 s). Dissociation-by-dilution experiments determined a k_off_ of 0.56 min^-1^ (95% CI: 0.48-0.67 min^-1^) corresponding to a half-life of 1.23 min (95% CI: 1.04-1.45 min), and a calculated k_on_ of 8.1 × 10^4^ M^-1^ min^-1^ (Supplementary Fig. 2e,f). The properties of TrimFABS_ANOX_ suggest that it is suitable for real-time monitoring of trimethoprim dynamic *in vivo* under oxic and anoxic conditions.

We tried to engineer an anoxic version of TetFABS that is anaologous to TrimFABS_ANOX_, yet these attempts were not successful. This is possibly due to differences in allosteric coupling of the sensory units and cpEGFP in the two cpEGFP-based biosensors. The fluorescent unit in TrimFABS is inserted in a flexible region of DHFR, which undergoes a major conformational change upon antibiotic binding (Fig. 1c), and which likely translates into altered environments around the chromophore irrespective of the nature of the fluorescent unit. In contrast, in TetFABS no such large-scale movements are predicted for the insertion site (Fig. 1h) suggesting that allosteric coupling is established by a much more specific, intricate network of contacts between the sensory unit, the fluorescent unit and/or the connecting linkers. Thus, simply swapping the fluorescent unit likely destroys allosteric coupling in the TetFABS biosensor.

### TrimFABS_ANOX_ reveals antibiotic transport dynamics in bacterial biofilms

To characterize the dynamics of antibiotic penetration into bacterial biofilms, we grew *Vibrio cholerae* cells constitutively producing TrimFABS_ANOX_ and mScarlet-I as a reference signal into biofilm colonies of different diameters (50–500 μm), using microfluidic chambers with a constant flow of fresh medium. When biofilms reached the desired size, we stained the biofilms with the fluorogen HMBR (5 μM) until saturation then applied 25 μM trimethoprim and imaged the antibiotic penetration using time-lapse confocal microscopy every 2 seconds (Fig. 3d). Images of the oxygen-dependent fluorescent protein mScarlet-I reveal that biofilms with diameters >100 μm primarily display fluorescence near the biofilm boundary, while the fluorescence fades out towards the biofilm center. This highlights the importance of oxygen-independent biosensors for biofilms (Fig. 3e, left column). After antibiotic addition, TrimFABS_ANOX_ fluorescence increased throughout the whole biofilm indicating that trimethoprim reached all the bacterial cells (Fig. 3e, central *vs*. right images).

While fluorescence of the TrimFABS_ANOX_ signal rapidly satured across different zones of biofilms with diameters smaller than 150 μm, penetration of the drug into the inner regions of larger biofilms was delayed (Fig. 3f). To quantify this delay in antibiotic penetration, we measured the time needed to reach half saturation in the center of the biofilm after trimethoprim reaches the periphery of the biofilm. The time at which the cells in the biofilm periphery display an increased TrimFABS_ANOX_ signal corresponds to the time when the antibiotic reaches the biofilm, because TrimFABS_ANOX_ responds to the antibiotic within a few seconds (Supplemetary Fig. 2f). These quantifications showed that drug penetration time monotonically increases with biofilm diameter, yet for all biofilm sizes the trimethoprim penetration time was <2 minutes (Fig. 3g). Images revealed that all cells in the biofilm displayed increased intracellular trimethoprim levels and that there were no pockets of unexposed cells after 2 min of exposure.

### Trimethoprim distribution during bacterial infection of human urothelium

We next sought to test the anoxic trimethoprim sensor in more complex environments, where spontaneous plasmid loss or copy number effects may influence the readout. The FAST-based biosensor TrimFABS_ANOX_ shows a single excitation peak but lacks an intrinsic ratiometric readout. To engineer a simple normalization reference, we used the Matryoshka approach^31^ by inserting cyan (sfmTurquoise2ox) or red (mScarlet-I) fluorescent proteins into the loop connecting the original FAST N- and C-termini. Purification and characterization of the resulting sensor constructs showed that it retained its main spectral properties while adding the intended excitation peaks for signal normalization (Fig. 4a-d).

**Fig. 4.**
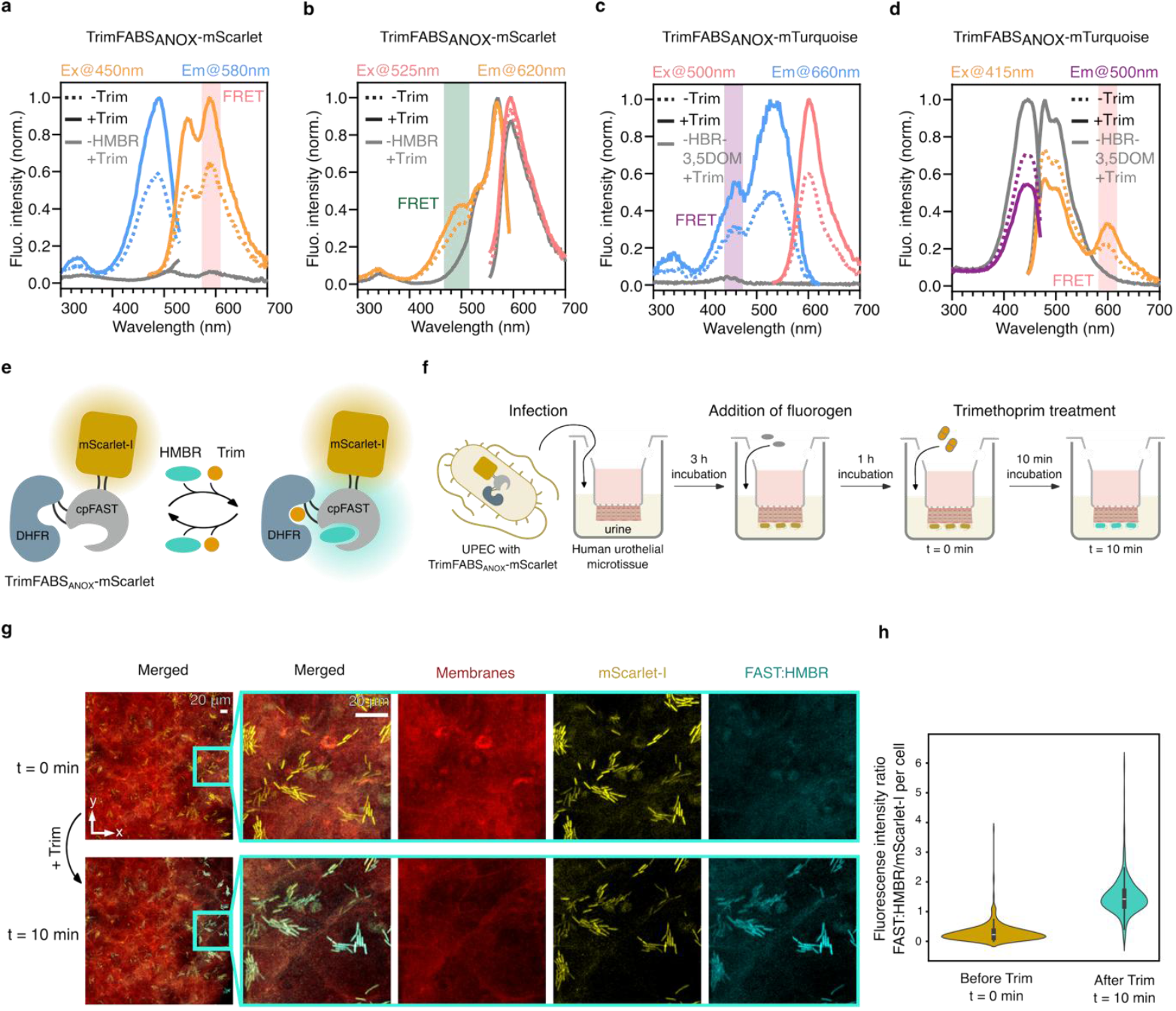
Matryoshka derivatives of TrimFABS_ANOX_ enable trimethoprim distribution quantification during UPEC infection of bladder tissue model. **a**, Excitation (blue) and emission (orange) spectra of TrimFABS_ANOX_-mScarlet in presence of 5 μM HMBR with and without Trim (solid/dashed lines, respectively). Adding only Trim without HMBR yields minimal fluorescence (grey). Excitation and emission wavelengths for emission and excitation spectra, respectively, are indicated on top and correspond to wavelengths that should capture FAST/HMBR-specific fluorescence, but not that of mScarlet-I; this is supported by the control sample lacking HMBR (grey lines), which shows only background fluorescence. The Trim-dependent extra peak at 590-600 nm (highlighted in red) results from Förster resonance energy transfer (FRET) between FAST/HMBR (emission maximum at ∼540 nm) and mScarlet-I (excitation maximum at ∼570 nm, with a second peak at ∼540 nm). **b**, Excitation (orange) and emission (red) spectra of TrimFABS_ANOX_-mScarlet analogous to panel a, except that the excitation spectra were measured at emission wavelength 620 nm, and the emission spectra were recorded using excitation at 525 nm. These wavelengths should specifically capture fluorescence of mScarlet-I, but only weakly the FAST:HMBR fluorescence. A green shaded region highlights FRET between FAST:HMBR and mScarlet-I moiety. **c**, Excitation (blue) and emission (red) spectra of TrimFABS_ANOX_-mTurquoise in presence of 5 μM HBR-3,5DOM with (solid) and without (dashed lines) Trim. Excitation and emission wavelengths for emission and excitation spectra, respectively, are indicated on top and correspond to wavelengths that should capture FAST/HBR-3,5DOM-specific fluorescence, but not that of sfmTurquoise2ox. **d**, Excitation (purple) and emission (orange) spectra of TrimFABS_ANOX_-mTurquoise in presence of 5 uM HBR-3,5DOM with (solid lines) and without (stippled lines) Trim, analogous to panel c, except that the excitation spectra were measured at emission wavelength 500 nm, and the emission spectra were recorded using excitation at 415 nm. **e**, Schematic representation of TrimFABS_ANOX_-mScarlet Matryoshka sensor. The sensory unit DHFR (blue) is coupled through linkers (black) with cpFAST (grey) and mScarlet (yellow) is linked to cpFAST providing reference fluorescence signal. TrimFABS_ANOX_ increases fluorescence upon simultaneous binding of trimethoprim (red) and a fluorogen of choice (for TrimFABS_ANOX_-mScarlet, we used HMBR). **f**, Schematic representation of the experiment: inverted human urothelial microtissue was infected with UPEC strain UTI89 harboring TrimFABS_ANOX_-mScarlet on a plasmid and incubated for 3 h. Then, urine as supplied with 5 μM HMBR. Images were taken with 40x objective before and 10 min after 100 μM trimethoprim addition. **g**, Top-view of the infected urothelial tissue with bacteria attached to the outside of the epithelial cells: before Trim (top panel) and after Trim (bottom panel) treatment: sensor (cyan), mScarlet (yellow), actin (red). **h**, Fluorescence intensity ratio between the FAST:HMBR channel and reference mScarlet channel before (yellow) and after (cyan) Trim addition, quantified per cell. All data points are represented as violin plots, where box plots show the median (white line), interquartile range (25th–75th percentiles; gray box), and whiskers extending to 1.5× the interquartile range.

Trimethoprim alone or in combination with sulfamethoxazole is routinely used to treat uncomplicated urinary tract infections caused by uropathogenic *Escherichia coli* (UPEC)^32,33^. Therefore, we applied the Matryoshka version of the trimethoprim sensor in a microtissue model of the human urothelium that displays an appropriate tissue stratification and cell differentiation, in which the apical side is exposed to human urine^34^. The urothelial tissue was infected with a UPEC strain producing the TrimFABS_ANOX_-mScarlet biosensor (Fig. 4e), treated with 100 μM trimethoprim and imaged before and after antibiotic addition in the same position (Fig. 4f). The concentration of trimethoprim we used is 100–400 times higher than the expected MIC^35^, which is within the typical range of trimethoprim concentrations detected in human urine during treatment^36^.

The resulting images revealed that trimethoprim reached all bacteria attached to the outside of the epithelial layer (Fig. 4g). The reference protein mScarlet enables the normalization of the fluorescence signal of the sensor by the protein copy number at the single-cell level by calculating the ratio between the fluorescent intensities of mScarlet and FAST:HMBR (Fig. 4h). These results demonstrate that TrimFABS_ANOX_-mScarlet is functional under highly physiological conditions using undiluted urine.

### Conclusion

This study presents, characterizes, and applies genetically encoded biosensors based on the activation of fluorescence by reversible allosteric binding of antibiotics (different tetracyclines or trimethoprim). These biosensors can be used to perform real-time fluorescence-based measurements of fluctuating concentrations of antibiotics in complex tissue environments, with high spatial and temporal resolution. Using these biosensors, we highlighted the important role of the host cell membrane in limiting tetracyclines transport, the short timescales of trimethoprim penetration into biofilms, the importance of oxygen-independent biosensors in biofilms, and the application of such sensors in a bladder tissue model. Antibiotic biosensors now enable detailed mechanistic studies of antibiotic transport across eukaryotic and prokaryotic cell membranes. Additionally, these biosensors can be applied in animal models and tissue models of infection, to reveal the impact of antibiotic pharmacokinetics and pharmacodynamics on treatment failure. The design strategy for antibiotic biosensors that we presented here can be employed for the urgently-needed development of biosensors for additional classes of clinically relevant antibiotics.

## Materials and Methods

### Culture conditions and medium

Prior to every experiment, overnight cultures of *E. coli, S. enterica*, or *V. cholerae* strains were grown in Lysogeny Broth (LB) Miller liquid medium (Roth) with shaking at 250 rpm. The incubation temperature was 37 °C for *E. coli* and *S. enterica*, and 28 °C for *V. cholerae*.

For biofilm assays in the microfluidic flow chambers, M9 medium was used for cultivation of *V. cholerae*. M9 medium contained 11.33 g L^-1^ Na_2_HPO_4_ · 7 H_2_O (Sigma, S9390-100G), 3 g L^-1^ KH_2_PO_4_ (Roth, 3904.2), 5 g L^-1^ NH_4_Cl (Roth, K298.1), 0.5 g L^-1^ NaCl (Roth, HN00.2), 0.492 g L^-1^ MgSO_4_ · 7 H_2_O (Roth, P027.1), 0.0111 g L^-1^ CaCl_2_ (Sigma Aldrich, C5670-100G), MEM vitamin Solution (Sigma, M6895-100ML), 2.78 g L^-1^ triethanolamine hydrochloride (pH 7.1) (Sigma, T1502-1KG), 0.00162 g L^-1^ FeCl_3_ (Roth, P742.1), 5 g L^-1^ D-(+)-glucose (Sigma, G8270-1KG), and trace elements (1.8 μg L^-1^ ZnSO4 · 7 H_2_O (Sigma-Aldrich, Z4750), 1.2 μg L^-1^ CuCl_2_ · 2 H_2_O (Roth, CN82.1), 1.2 μg L^-1^ MnSO_4_ · H_2_O (Sigma, M7634-100G), 1.8 μg L^-1^ CoCl_2_ · 6 H_2_O (Sigma, C8661-25G)).

Where required, the culture medium was supplemented with the following antibiotics: gentamicin (30 μg mL^-1^), ampicillin (250 μg mL^-1^), or kanamycin (50 μg mL^-1^). The concentrations of tetracyclines and trimethoprim varied between assays and are listed in the respective figures.

### Plasmid construction

Plasmid construction was carried out using standard molecular biology techniques. All enzymes for cloning were purchased from New England Biolabs (NEB), if not specified otherwise. DNA purification from gels were carried out with Macherey-Nagel NucleoSpin Gel and PCR Clean-up. Primers were designed using the SnapGene software (Insightful Science) or Geneious Prime (Biomatters Ltd). Oligos used for plasmid construction were synthesized by Sigma Aldrich or Microsynth (Balgach, Switzerland). Synthetic genes were ordered from Twist Bioscience. All constructed plasmids were purified with Macherey-Nagel NucleoSpin Plasmid Easy-pure kit or GenElute Miniprep kit (Sigma-Aldrich) and verified by Sanger sequencing, which was carried out by Eurofins or Microsynth. The plasmid pUC-Mu2 was additionally subjected to FullPlasmidSeq by Microsynth.

### Bacterial strain construction

Electroporation (MicroPulser, Bio-Rad) was used to deliver pooled plasmid libraries into the *E. coli* MegaX DH10B T1 strain. Electroporation was also used for plasmid transformation into UPEC strains UTI89 and CFT073, as well as *S. enterica*. After electroporation, the cell suspension was supplemented with SOC medium and recovered for 40 minutes at 37 °C. *E. coli* BL21-Star, Top10, and S17 strains were transformed via heat-shock, using a Gibson reaction or ligation product. *V. cholerae* strains were transformed by conjugation with *E. coli* S17-λ pir harboring the desired plasmid.

### Construction of Mu transposon-based domain-insertion biosensor libraries

Biosensors were constructed by implementing a method by Nadler et al.^16^ based on creating random domain-insertion libraries, with modifications as outlined in the following. Plasmid pConRef-DHFR (pNUT4499) harboring the DHFR sensory unit was constructed via Gibson combining linearized with oligos kdo5127 and kdo5128 pConRef-2H12 and synthetic codon-optimized sensory unit sequence with addition of RBS amplified with oligos kdo5116 and kdo5117. Plasmid pCon-TetR harboring the TetR sensory unit from a constitutive promoter was constructed by PCR amplification of *tetR* from p2H12^37^ with oligos 18995/18996 and cloning in pConRef-2H12 digested with EcoRI/NheI via Gibson assembly, replacing the *cdGreen*-*mScarlet-I* fragment.

For *in vitro* transposition reactions, a modified mini-Mu transposon containing a kanamycin resistance cassette was constructed by PCR amplification of *nptII* from pAK405^38^ using oligos 12292/12293, digestion with HindIII and ligation in HindIII-cut pUC18. The resulting plasmid, called pUC-Mu2, releases transposition-competent Mu transposon (Mu2) upon HindIII digestion and harbors the same BsaI-containing Mu-ends described by Nadler et al^16^.

*In vitro* transposition reactions were carried out in 20 μL reaction containing 150 ng plasmid, 100 ng HindIII-released and gel-purified Mu2 transposon, 1x MuA buffer, and 0.22 μg of MuA transposase enzyme (Thermo Fisher Scientific). Reactions were incubated at 30 °C for 18 hours followed by 15 min at 75 °C to inactivate the enzyme, purification using a PCR clean-up kit (Macherey-Nagel) with elution in MilliQ water, and transformation into *E. coli* MegaX DH10B T1 electrocompetent cells (Invitrogen). Following incubation for 1-2 hours in pure LB or SOC medium, transformations were transferred into at least 100 mL of liquid LB supplemented with ampicillin (250 μg mL^-1^) or carbenicillin (100 μg mL^-1^) and kanamycin (50 μg mL^-1^). An aliquot was plated on LB plates supplemented with the same antibiotics to assess the efficiency of the transformation and assure at least 5x coverage of possible Mu2 transposon insertion variants. Cultures were incubated for 16 h at 37 °C, 200-250 rpm. Plasmid library DNA was extracted with a plasmid Miniprep kit (Macherey-Nagel or Sigma-Aldrich) and eluted with MilliQ water.

To enrich the transposon-insertion libraries for Mu2 insertions inside the part encoding the sensory unit, the sensory unit part was excised from plasmid libraries using EcoRI/SpeI (for the trimethoprim sensor library) or EcoRI/NheI (for the tetracyclines sensor library) and ligated into clean backbones prepared with the same enzymes from the parental, unmutagenized plasmids. Ligations were then transformed into *E. coli* MegaX DH10B T1 electrocompetent cells (Invitrogen) and processed as described above for primary libraries. The coverage for both libraries were at least 100x. To exchange the Mu2 transposon with the *cpEGFP* gene, Golden Gate cloning was used as described in the following.

For trimethoprim sensor library, *cpEGFP*^16^ was PCR-amplified with kdo5170 and kdo5143 and purified from the gel. Golden Gate assembly was performed in total of 50 μL as follows: 500 ng Mu-insertion library, 500 ng gel-purified *cpEGFP*, 2.5 μL 400 U/μL T4 DNA ligase (NEB), 5 μL 10x T4 ligation buffer, 1.5 μL 20 U/μL BsaI-HFv2 enzyme (NEB). Reactions were incubated at 37 °C overnight followed by 15 min at 80 °C for enzymes inactivation. Resulting reaction products were purified, eluted in 30 μL of milliQ water, transformed into 50 μL of *E. coli* MegaX DH10B T1 electrocompetent cells (Invitrogen), and transferred into liquid LB medium supplemented with ampicillin (250 μg mL^-1^). An aliquot was plated on LB plates supplemented with the same antibiotic to assess the efficiency of the transformation. The culture was incubated for 16 h at 37 °C, with shaking at 250 rpm. The resulting plasmid library is termed “domain-insertion library” and comprises plasmids with randomly located insertions of *cpEGFP* in the gene coding for the sensory unit. Aliquots of the domain-insertion library were stored at -80 °C.

For the tetracyclines sensor library, *cpEGFP*^16^ or *cpEGFP*\*^37^ were PCR-amplified with oligo set 1 (equimolar mix of oligos 19376-19383) from template pTKEI-Tre-C04 (Addgene plasmid # 79754) or oligo set 2 (equimolar mix of oligos 19376-19379 and 19384-19387) from template p2H12 (Addgene plasmid # 221142), respectively, and PCR products were gel-purified. Golden Gate assembly was performed in a total of 50 μL containing 350 ng of pCon-TetR Mu-insertion library, 250 ng of *cpEGFP* or *cpEGFP*\*, 2.5 μL 400 U/μL T4 DNA ligase (NEB), 1.5 μL 20 U/μL BsaI-HFv2 enzyme (NEB), and 5 μL 10x T4 ligase buffer. Golden Gate assembly reaction conditions and following steps were as described above, except that carbenicillin (100 μg mL^-1^) was used for selection.

### Fluorescence-activated cell sorting (FACS) of trimethoprim biosensor variant libraries

Several iterations of fluorescence-activated cell sorting (FACS) were performed to identify functional biosensors in the *cpEGFP*-insertion libraries. Libraries were grown overnight in LB containing ampicillin (250 μg mL^-1^) at 37 °C with shaking at 250 rpm, then diluted 1:100 in the fresh LB with ampicillin (250 μg mL^-1^) and grown at 37 °C with 250 rpm until OD_600_ ∼ 0.4-0.6. Afterwards, cultures were diluted 1:100 in phosphate-buffered saline (PBS), split into two equal volumes and placed on ice. 10 μM trimethoprim was added to one of the two tubes, and both tubes were incubated on ice for 15 minutes. FACS was performed on the Aria III cell sorter (BD Biosciences) utilizing a 488 nm excitation laser with a 514/30 emission filter for GFP and a 561 nm excitation laser with a 610/20 emission filter for mScarlet. For analysis and setting the gates, 10,000 events were recorded. Afterwards, 100,000 cells were sorted into 1 mL of LB, incubated for 1 h at 37 °C with shaking at 750 rpm, and then transferred into 3 mL of LB supplemented with 250 μg mL^-1^ ampicillin for overnight incubation. The overnight culture was then used for the next iteration of the same FACS procedure described above to further enrich for functional biosensors, or stored as a glycerol stock at -80 °C.

The gates in each round of FACS were designed to selecting for the GFP-positive cell population in the sample with antibiotics added, and for the GFP-negative cell population without antibiotics. Several iterations of FACS followed by overnight growth of the sorted populations were carried out until a clear difference in GFP fluorescence intensity between the antibiotic-treated and antibiotic-untreated cells was detected. The resulting library was plated on LB agar plate supplemented with 250 μg mL^-1^ ampicillin to select single colonies containing one variant of the biosensor. For each selected variant of the biosensor, the change of the fluorescence in absence *vs*. presence of the antibiotic was then evaluated with a Fortessa cell analyzer (BD Biosciences, 488 nm excitation laser with 512/25 emission filter). The resulting data was visualized with FlowJo. For those biosensor variants for which the Fortessa cell analyzer confirmed a fluorescence shift, the strain harboring the biosensor candidate was cultured overnight in LB supplemented with 250 μg mL^-1^ ampicillin, and DNA was isolated and sequenced with Sanger sequencing.

To improve brightness and response amplitude of the biosensor candidates that were obtained from the iterative FACS procedure described above, one round of optimization was carried out. *cpEGFP* was amplified with randomized linker sequences (oligos: kdo5332, kdo5294) and assembled via Golden Gate with the pConRef plasmid carrying the sensory unit linearized with primers kdo5292 and kdo5293. The resulting library was purified, eluted in 30 μL of milliQ water, transformed into 50 μL of MegaX DH10B T1 electrocompetent cells (Invitrogen) and transferred into liquid LB medium supplemented with ampicillin (250 μg mL^-1^). An aliquot was plated on LB plates supplemented with the same antibiotics to assess the efficiency of the transformation. The culture was incubated for 16 h at 37 °C with shaking at 250 rpm. Aliquots of the library containing the randomized linker sequences for *cpEGFP* were stored at -80 °C or used directly for another round of the iterative FACS procedure as described above, to select variants in which the cpEGFP linker improved the dynamic range of the biosensor candidates. The resulting biosensor variant we called TrimFABS (encoded on the plasmid pConRef-Trim2.2) and characterized in detail in this study.

### FACS-based isolation of first- and second-generation tetracycline biosensors

For FACS of TetR domain-insertion libraries, cells were grown overnight in 5 ml LB containing carbenicillin (100 μg mL^-1^) at 37 °C with shaking at 200 rpm. 200 μL of the overnight culture were then diluted in 5 mL of PBS with or without 200 nM anhydrotetracycline (aTc), incubated for 1 h at room temperature, and then subjected to FACS on an Aria III sorter (BD Biosciences) using a 488 nm excitation laser and a 514/30 emission filter for GFP. After a first sort for GFP^+^ cells in presence of aTc (defined by a NOT-gate set by a negative gate with non-fluorescent bacteria), subsequent FACS rounds were performed with alternating selection regimes, sorting for the top or bottom 2-5% in the presence or absence of aTc, respectively. In each sorting round, sorted cells (typically >50’000) were directly collected in 5 ml LB containing carbenicillin (100 μg mL^-1^) and then incubated at 37 °C with shaking at 200 rpm. In case the sort was before a weekend, the culture was left at room temperature during the weekend, was diluted back 1:10 on Monday and incubated for 1 h at 37 °C with shaking before being diluted in PBS plus/minus aTc and processed as described above. For first-generation biosensors, a total of six FACS rounds were applied before single colonies were being isolated and tested individually.

To test individual clones for their response to aTc, an aliquot of the overnight culture resulting from the final FACS selection was streaked on an LB plate containing carbenicillin (100 μg mL^-1^), incubated overnight at 37 °C, and individual colonies were picked and inoculated into 100 μL of LB containing carbenicillin (100 μg mL^-1^) in 96-well U-bottom tissue culture plates (Falcon; Cat# 353077) and grown in an Synergy H1 plate reader overnight at 37 °C with double-orbital shaking at max speed. Cells were pelleted by centrifugation of the 96-well plates in a model 5810 R centrifuge (Eppendorf) at 300 rpm, 10 min, room temperature. Supernatants were carefully aspired, and pellets resuspended in 145 μL of PBS, followed by recording of GFP fluorescence (excitation: 485 nm, emission: 528 nm) and absorbance at 600 nm (OD_600_) values at 37 °C every 3 min for 10 min. Then, 5 μL of a 33.3x stock of aTc in PBS were added to each well and recording of GFP and OD_600_ values was continued for 1 h. Fluorescence values were normalized by the OD_600_ and fold changes were calculated for each time point after aTc addition relative to the last time point before aTc addition. For a total of 16 clones based on the largest fold changes plasmids were isolated and Sanger sequenced, and one of these variants (encoded on plasmid pCon-TetR5), was chosen for further rounds of linker optimization.

For linker optimization, *cpEGFP*\* was amplified from pCon-TetR5 in four separate PCR reactions containing forward oligo 20192 and one each of reverse oligos 20195-20198. The plasmid backbone was linearized by PCR using oligos 20188/20189 and pCon-TetR5 as template. All PCR reactions were gel-purified and used in a single Golden Gate assembly reaction containing 140 ng of backbone PCR and approximately 130 ng of each of the four *cpEGFP\** PCR reactions, 2.5 μL 400 U/μL T4 DNA ligase (NEB), 1.5 μL 20 U/μL BsaI-HFv2 enzyme (NEB), and 5 μL 10x T4 ligase buffer. Golden Gate assembly reactions, subsequent transformations and iterative FACS were essentially performed as for first-generation tetracycline sensors, except that the first round of selection for GFP^+^ cells was omitted due to a clear aTc dependency already of the naïve linker library. After five rounds of FACS, single clones were tested individually in 96-well format as described above, and sensor plasmids from the twelve best-performing clones were isolated and sequenced, revealing a single unique variant (encoded on plasmid pCon-TetR5.7) that we further characterized as our final tetracycline sensor.

### Synthesis of HMBR and HBR-3,5-DOM fluorophores

HMBR was synthesized as previously described in Plamont et al., 2016^29^: A solution containing rhodanine (202 mg, 1.52 mmol) (Thermo Scientific) and the 4-hydroxybenzaldehyde (195 mg, 1.60 mmol) (Sigma) in 110 mL of water was stirred at 65-80 °C for 10 days. After cooling to room temperature, the precipitate was filtered through a glass filter, washed with water, and dried under vacuum. HMBR was obtained as a dark yellow powder (45%).

HBR 3,5-DOM was synthesized as previously described in Li et al., 2017^30^: A solution containing rhodanine (1.13 mmol, 1.0 eq) (Thermo Scientific), 4-hydroxy-3,5-dimethoxybenzaldehyde (1.24 mmol, 1.1 eq) (Apollo) and piperidine (1.13 mmol, 1.0 eq) in 5.5 mL absolute ethanol was stirred at room temperature (28 °C) for 48 h, before addition of water (20 mL) and neutralisation with hydrochloric acid (dropwise, 10% (v/v)). The solid was washed on a glass filter with water and absolute ethanol and dried under vacuum. HBR 3,5-DOM was obtained as a brown powder (33%).

### FACS-based selection of cpFAST-based trimethoprim biosensor

To convert the GFP-based trimethoprim biosensor into a FAST-based biosensor, synthesized codon-optimized circularly permutated *FAST* (*cpFAST*) was PCR-amplified with a primer pool encoding randomized linkers with varying lengths (oligos: kdo5442-5446) and cloned into the same position of *DHFR* where *cpEGFP* was located in pConRef-Trim2.2 (oligos for PCR: kdo5292, kdo5293) via Golden Gate assembly as described before. Libraries with different linker length were pooled together and aliquots of the pooled stored at -80 °C or used directly for the iterative FACS. Before the FACS, libraries were grown overnight in LB containing ampicillin (250 μg mL^-1^) at 37 °C with shaking at 250 rpm, then diluted 1:100 in the fresh LB with ampicillin (250 μg mL^-1^) and grown at 37 °C with 250 rpm until OD_600_ ∼ 0.4-0.6. Afterwards, cultures were diluted 1:100 in phosphate-buffered saline (PBS), split into two equal volumes, incubated with 5 μM HMBR (^TF^Lime, Twinkle Biosciences or self-made) fluorophore for 1 hour at room temperature. After that, samples were placed on ice and 10 μM trimethoprim was added to one of the two tubes for 15 min. The iterative FACS and further single colony screening to select the final cpFAST-based biosensor was identical to the selection of the cpEGFP-based trimethoprim biosensor. The resulting biosensor was called TrimFABSANOX (encoded on the plasmid pConRef-Trim-FAST) and characterized in this work.

### Protein purification

Biosensors were cloned in pET28a-based plasmids for production of amino-terminally His6-tagged proteins. For construction of pET28-TetR5.7, a fragment harboring *tetR5*.*7* was released from pCon-TetR5.7 by digestions with EcoRI/NheI and ligated in EcoRI/SpeI-digested pET28-2H12, replacing *cdGreen*. pNUT4134 (pET28 with TrimFABS) was constructed as follows: TrimFABS sequence was amplified with kdo5622 and kdo5512 from pNUT3797, digested with EcoRI/NheI and ligated in EcoRI/SpeI-digested pET28-2H12, replacing *cdGreen*. For construction of pET28-Trim-cpFAST-MatryScar and pET28-Trim-cpFAST-MatryTq, *E. coli* codon-optimized synthetic genes were ordered from TWIST Bioscience, digested with EcoRI/NheI, and cloned in EcoRI/SpeI-digested pET28-2H12. pET28-Trim-cpFAST was constructed by release of *mScarlet-I* from pET28-Trim-cpFAST-MatryScar by KpnI digestion, followed by religation of the backbone.

Biosensors were overexpressed in *Escherichia coli* Lemo21(DE3). Cells were grown at 37 °C in Terrific Broth medium containing 50 μg mL^−1^ kanamycin to an OD_600_ of 0.8–1.0. Protein production was induced with 0.1 mM IPTG, then the temperature was reduced to 20 °C and cells were harvested after overnight incubation. Cells were lysed with a sonicator in 50 mM Tris-HCl pH 8.0, 250 mM NaCl, 40 mM imidazole, and the soluble protein was purified by immobilized metal affinity chromatography (IMAC) on a Ni Sepharose column HP column (Cytiva) and subjected to size-exclusion chromatography on a Superdex S200 column (Cytiva) in sensor buffer (25 mM Tris-HCl pH 7.5, 150 mM NaCl, 10 mM MgCl_2_, 10 mM KCl, 5 mM β-mercaptoethanol). Protein-containing fractions were pooled and concentrated in Amicon Ultra units (Merck). Protein yields were 10-20 mg L^−1^. Purified biosensors were aliquoted and stocked at −80 °C for long-term storage.

### *In vitro* characterization of biosensors

All biosensor measurements were performed on SynergyH4 or SynergyH1 plate readers (BioTek Agilent) in a total volume of 100-200 μl sensor buffer (25 mM Tris-HCl pH 7.5, 150 mM NaCl, 10 mM MgCl_2_, 10 mM KCl, 5 mM β-mercaptoethanol) at 30°C using 96-well polystyrene tissue culture-treated flat bottom plates (Falcon).

For dose-response curves and spectral scans, reactions contained 250 nM biosensor and ligands and cofactors at concentrations indicated in the figures or figure legends prepared from the following fresh stock solutions: 10 mM anhydrotetracycline hydrochloride (Fluka) in DMSO; 20 mM tetracycline hydrochloride (Carl Roth) in methanol; 20 mM oxytetracycline hydrochloride (Sigma) in ddH_2_O; 45 mM chlortetracycline hydrochloride (Sigma) in ddH_2_O; 40 mM doxycycline hyclate (Sigma) in methanol; 44 mM minocycline hydrochloride (Sigma) in 75% methanol; 50 mM trimethoprim (Acros Organics) in DMSO; 14 mM NADPH tetrasodium salt (Sigma) in ddH_2_O; 5 mM HMBR (in-house synthesized, or purchased as ^TF^Lime from The Twinkle Factory), HBR-3,5DM (^TF^Amber from The Twinkle Factory) and HBR-3,5DOM (in-house synthesized, or purchased as ^TF^Coral from The Twinkle Factory) in DMSO. Biosensors with ligands and cofactors were typically equilibrated at 30°C for 1h prior to measurements. For trimethoprim biosensor dose-response curves in the absence and presence of one of the ligands/cofactors (NADPH, trimethoprim and/or HBMR), dose-response curves were first recorded without the individual ligand/cofactor, then the ligand/cofactor was added from a concentrated stock solution (<10%[vol/vol] of total assay volume), reactions were equilibrated for an additional hour, and dose-response curves were recorded again. Multiple data points were determined for each sample using repeated measurements of the same well every 1-10 min over a period of up to 7h, and later data points at which binding was in equilibrium were used for dose-response curves. K_d_ values were fitted using GraphPad Prism 10 software and the “[Inhibitor] vs. response - Variable slope (four parameters)” model. For spectral scans, saturating ligand concentrations were used (1.25 mM trimethoprim, 20 μM anydrotetracycline, 250 μM NADPH, 5 μM fluorogen) and excitation and emission spectra were recorded with 1-2 nm resolution. For TrimFABS excitation spectra shown in Fig. 1d, spectra of NADPH alone were subtracted from the recorded biosensor spectra since NADPH itself shows a major excitation peak around 300-400 nm.

For “dissociation by dilution” experiments to determine k_off_ rate constants, the following mixes were prepared in 20-50 μl sensor buffer: 2.5 μM TrimFABS, 250 μM NADPH and 10 μM trimethoprim; TrimFABS_ANOX_, 5 μM HMBR and 10 μM trimethoprim; or 100 μM TetFABS and 20 μM doxycycline. After equilibration for 1h at 30°C, mixes were diluted 75-100-fold in buffer without antibiotics and biosensor, but with other cofactors, and measurements were started immediately (ca. 10 s dead time). k_off_ values were fitted using GraphPad Prism 10 software and the “Dissociation - One phase exponential decay” model. Association kinetics were measured by incubation of 250 nM biosensors (with appropriate cofactors as required) in the absence of antibiotic ligands in sensor buffer for 1h, followed by addition of antibiotics from concentrated stock solutions to saturating concentrations (20 μM doxycycline or 500 μM trimethoprim final concentrations) and immediately starting recording of biosensor responses (ca. 10 s dead time). Saturation half-time was defined as the time needed to reach 50% of the maximal fluorescent biosensor signal and calculated from the mean of triplicates.

### Macrophage infection assays with *S*. *enterica*

TrimFABS and TetFABS were subcloned into pNUT542 instead of double GFP, excluding mKOKappa protein and exchanging gentamycin resistance to kanamycin resistance. For the first Gibson assembly, backbone of pNUT542 was PCR amplified with oligos kdo5428 and kdo5429, promotor region and RBS were amplified from pNUT542 with kdo5430 and 5431 and TrimFABS was amplified from pConRef-Trim2.2 with oligos kdo5432 and kdo5433. The resulting plasmid pNUT4096 was amplified with kd70 and kdo1114 leaving out gentamycin resistance gene and assembled via Gibson with kanamycin resistance fragment amplified with kdo1773 and kdo851. The resulting pNUT4455 carrying TrimFABS sensor was amplified with kdo5469 and kdo5470 to exchange TrimFABS with TetFABS amplified with kdo5434 and kdo5435 from pConRef-TetR5.7. PCR fragments were assembled via Gibson and resulted in plasmid pNUT4394. The resulting plasmids were sequenced with Sanger sequencing and electroporated into kds016.

The macrophage cell line RAW 264.7 was used for these assays. Macrophages (3.5 × 105/well) were added to 12-well glass-bottom plates (Cellvis) and supplemented with DMEM cell culture medium (Gibco, 21969-035) containing 10% fetal bovine serum (Gibco, 10270-106) and 1% GlutaMax (Gibco, 35050038) for 24 hours prior to bacterial infection.

Macrophages were infected with the *Salmonella* SL1344 wild-type strain producing either the TetFABS (from pNUT4394) or TrimFABS (from pNUT4455) sensor at a MOI of 10:1. The infected cells were then centrifuged at 400 g for 5 min to synchronize internalization. After 30 min, macrophages were washed with media containing 50 μg/mL gentamicin to remove non-internalized *Salmonella*. Fresh media containing 50 μg/mL gentamicin was then added to the cells for the duration of the infection.

At 4 hours post-infection, macrophages were washed and stained with CellTracker Deep Red Dye (ThermoFisher, C34565; final concentration 3 μM) and Hoechst 3342 (ThermoFisher, 62249; final concentration 20 μM; suspended in cell media) for 20 min at 37 °C in a humidified 5% CO_2_ incubator. Stained macrophages were washed three times in cell media to remove the staining solution, and fresh media containing 50 μg/mL gentamicin was then added back to the cells for the duration of imaging.

On the microscope, cells were incubated at 37 °C and in an atmosphere containing 5% CO_2_. Fluorescence was imaged in 4 channels (GFP, mScarlet, DAPI, Deep red) every 20 s (TrimFABS) or 60 s (TetFABS). First, 3-6 fields of view at a time were imaged for 2 minutes to record the baseline signal. After 2 minutes, equal volume of DMEM cell culture medium with 100 μM trimethoprim or 100 μM doxycycline with or without 0.2% Triton X-100 (Sigma, T8787-50ML) was added to each well. Fluorescence was imaged in 4 channels (GFP, mScarlet, DAPI, Deep red) every 20 s (TrimFABS) or 60 s (TetFABS). Imaging was stopped after 30 – 60 min.

Imaging was performed with a Yokogawa CSU-W1 confocal spinning disk unit mounted on a Nikon Ti-E inverted microscope using a 40x oil objective with numerical aperture 1.3 (Nikon) or 100x oil objective with numerical aperture 1.45 (Nikon). Fluorescent proteins harbored by *Salmonella enterica* were excited with a 488 nm laser (TrimFABS, TetFABS) or a 552 nm laser (mScarlet), and macrophages were visualized by exciting the applied dyes with a 637 nm laser (CellTracker Deep Red Dye) and a 405 nm laser (Hoechst 3342). The microscope hardware was controlled by NIS Elements (Nikon). Images were captured by an Andor iXon 888 Ultra EMCCD camera, cooled to -70 °C, in one z-plane among several xy positions. A Nikon hardware autofocus was used to correct focus drift.

Bacteria culture without macrophages controls were obtained as follows: *Salmonella* SL1344 wild-type strain producing either the TetFABS (from pNUT4394) or TrimFABS (from pNUT4455) sensor were grown overnight in LB containing 50 μg/mL kanamycin, diluted 1:100 in 2 ml of cell culture medium in 12-well plate and incubated for 4 h at 37 °C in a humidified 5% CO_2_ incubator. After incubation, 2 ml of bacterial suspension were centrifuged at 6000 g for 2 min and resuspended in 1.5 ml of PBS. In 96-well flat-bottom plate (Falcon), 100 μl of bacterial suspension in PBS were mixed with 100 μl of 100 μM doxycycline, 100 μM tetracycline or pure PBS with and without 0.2% triton and immediately put into the plate reader (BioTek, Synergy H1 microplate reader). Fluorescence intensity was measured for 1 h every 30 sec for mScarlet and GFP. All measurements were done in technical triplicates.

### Image analysis of antibiotic penetration in macrophage infections

Imaging data acquired with 40x oil objective was analyzed using the BiofilmQ software^39^. To quantify the fluorescence increase of the biosensors over time, we analyzed two imaging channels: mScarlet as a constitutive signal and GFP as the sensor signal. Cell segmentation was performed on the constitutive signal, using Otsu’s method for threshold calculation with a manually adjusted sensitivity. Before the segmentation of the bacterial cells the raw images were denoised by convolution with an averaging kernel (size corresponds to default values of BiofilmQ) and top-hat filtering with a disk-shaped kernel slightly larger than the expected bacterial size (here 2.7 μm or 7 px). Segmented clusters smaller than 0.10 μm^3^ were subsequently removed. Time-resolved GFP and mScarlet fluorescence intensity per segmented object was calculated extracting the fluorescence values from the raw images on the segmented objects. The following set of BiofilmQ-parameters were computed, exported and plotted into graphs: “Global biofilm properties” and “Fluorescent properties”, namely “Integrated intensity per object” in every channel and “Integrated intensity ratio per object” (ratio between mScarlet and GFP fluorescence intensities with background subtraction). Half-saturation time (T_50_) was defined as the time when normalized fluorescence intensity ratios reached 50% of the maximum. Statistical significances of the half-saturation times were calculated between triton-treated and untreated conditions using a two-way Mann-Whitney-Wilcoxon test.

For bacterial suspension controls without macrophages, fluorescence intensity values were directly taken from the plate reader. Then we calculated the ratio between sensor (GFP) fluorescence intensity and mScarlet. Time between mixing and the first plate reader measurement was ∼30 sec, therefore, fluorescence in the wells without antibiotics at the first time point was averaged and taken as intensity at 0 sec. Half-saturation time (T_50_) was defined and plotted as listed above.

### Antibiotic penetration assays of *V*. *cholerae* biofilms

For microfluidic flow chamber construction, the polydimethylsiloxane (PDMS) and glass coverslips were bonded using an oxygen plasma, resulting in flow chambers that were 10 mm in length, 4 mm in width, and 500 μm in height. The microfluidic design contained 7 independent channels of identical dimensions, each with its own inlet and outlet, on a single chip.

Plasmid pNUT4098 harboring TrimFABS_ANOX_ was created via Gibson from pNUT4096 by substituting TrimFABS. Backbone from pNUT4096 was PCR amplified with oligos kdo5428 and kdo5431 and pConRef-Trim-FAST (pNUT4086) was PCR amplified with oligos kdo5432 and kdo5433. The resulting plasmid was sequenced with Sanger sequencing.

For biofilm assays, a *V. cholerae* strain producing the FAST-based trimethoprim biosensor (TrimFABS_ANOX_) was created via delivering the plasmid pNUT4098, which contains a gentamicin resistance marker, into strain kdv3820. Strain kdv3820 is a derivative of the N16961 wild type strain carrying the following mutations: *vpvC*^W240R^ (rugose allele), Δ*flaA*, Δ*mshA*. The resulting *V. cholerae* strain (kdv3533), which constitutively produced TrimFABS_ANOX_ and mScarlet, was cultured at 28 °C with shaking at 250 rpm, in liquid LB containing 30 μg/mL gentamicin to maintain the plasmid pNUT4098. After 16 h of incubation, the culture was diluted 1:100 in fresh medium and cultured in the same conditions until OD_600_ = 0.4. At this point, the culture was diluted 1:600,000 in LB without antibiotics and inoculated into the microfluidic flow chamber, which resulted in approximately 1-2 biofilm colonies per channel. After inoculation, the bacterial suspension was incubated in the microfluidic chambers for 1 hour without flow, to allow *V. cholerae* attachment to the glass surface. Then, every channel was connected to a syringe pump using tubing with inner diameter of 0.3 mm (MasterFlex, MFLX06417-11). Unattached bacterial cells were then washed out of the flow channels by a 1-min long pulse of strong flow (100 μL/min) of sterile, freshly prepared M9 medium. Afterwards, the flow rate of sterile M9 medium was reduced to 10 μL/min and biofilms were grown at 25 °C for 20-60 hours to result in biofilm colonies with diameters between 50 and 400 μm.

After biofilms reached the desired size, the inflowing medium was exchanged from simple M9 medium to M9 medium containing 5 μM HMBR (^TF^Lime) for 1-2 hours. Then, the inflowing medium was exchanged from M9 medium containing 5 μM HMBR to M9 medium containing 5 μM HMBR and 25 μM trimethoprim. The media exchanges were performed using a microfluidic connector chip, which enabled us to switch the medium that flowed into the flow channel without interrupting the flow. The microfluidic chips were imaged using the same confocal microscope that was used to image the macrophage infection assays (described above), but for the biofilm assays we used a 20x air objective with numerical aperture 0.75 (Nikon). After the medium exchange to medium containing HMBR, images were acquired in one z-plane at the bottom of the biofilm every 2 seconds in two fluorescent channels, excited by 488 nm laser (TrimFABS_ANOX_) and a 552 nm laser (mScarlet).

### Image analysis of antibiotic penetration in biofilms

For each timepoint in the timeseries, the 2D image was segmented into biofilm *vs*. background based on the TrimFABS_ANOX_ signal. Segmentation was performed via Otsu^40^ thresholding from the scikit-image library^41^. All pixels with values bigger than 0.9x the Otsu threshold were set to foreground pixels. In a subsequent step, only the largest connected component was determined as the biofilm. This segmentation mask was saved and inspected visually. In cases where the segmentation was erroneous, the respective segmentation was curated manually.

To add spatial context to the segmentation, the Euclidean distance transform - which approximates the distance to biofilm boundary for each pixel - of each segmentation in the timeseries was calculated^42^ based on the pixel size. Next, these distances were digitized into 100 bins. For each distance bin, mean and standard deviation of the intensities of all pixels belonging to that bin were calculated. Giving rise to a spatio-temporal kymograph of TrimFABS_ANOX_ intensities, that subsequently was smoothened by calculating the running mean with a kernel size of five timesteps along the temporal axis. To summarize the intensity dynamics visible in the kymograph, we approximated the half penetration time^43^. Penetration time is the time period from the very start of the penetration of a molecule into the biofilm until the timepoint the center of the biofilm reaches the concentration present in the medium. As concentration changes lead to changes in intensities, we defined penetration time as the period from the 5% of signal increase in the most peripheral biofilm bins until TrimFABS_ANOX_ intensity in the center bins increased to 50% of the maximal value similar to the half-saturation time (T_50_) above.

### Infection assays of human 3D urothelium microtissue model

Plasmid pNUT4211 harboring TrimFABS_ANOX_-mScarlet was created via Gibson from pNUT4096 by substituting TrimFABS. pNUT4096 was PCR amplified with oligos kdo5633 and kdo5428 and pET28-Trim-cpFAST-Matry-Scar PCR amplified with oligos kdo5631 and kdo5632. The resulting plasmid pNUT4211 was partially sequenced with Sanger sequencing and electroporated into the UTI89 strain kde3333 resulting in kde4498 strain.

Urine-tolerant human urothelial tissues were generated on the underside of 24-well transwell inserts as described before^34^. In brief, HBLAK cells (passage 10–12) were seeded onto 6.5-mm polycarbonate transwell inserts with 0.4-μm pore size (Corning 3413) at a density of 2.5×106 cells mL^−1^ using an inverted seeding method and incubated at 37 °C with 5% CO_2_. After 4 h, 2D cultivation medium (CnT-Prime, CELLnTEC) was added. After 16 h, the inserts were returned to normal orientation, and 2D cultivation medium was added to both chambers (day 1). To validate tissue development, Invitrogen CellMask Deep Red Plasma Membrane Stain (C10046, Thermo Fisher, Waltham, MA) was added to the medium (dilution 1:5000) Confluence was reached by day 4, at which point medium was replaced with 3D-differentiation medium (CnT-PR-3D, CELLnTEC) supplemented with a growth factor mix (00192152, KGM Gold SingleQuots, Lonza); no antibiotics were used. Medium was refreshed on days 6, 8, 11, and 13. On day 15, cultures were placed at an air-liquid interface and maintained until day 22 in unsupplemented 3D differentiation medium. On day 22, a urine-liquid interface was established by adding pooled human urine to the lower chamber. 3D differentiation medium and urine were exchanged every 2 days, and experiments were performed on days 24-26.

For the pooled human urine, urine was collected anonymously from healthy volunteers of both sexes, without diabetes mellitus, antibiotic treatment within the previous 14 days, known pregnancy, menstrual bleeding on the day of sample collection or current urinary tract infection. Individual urine samples were analyzed using a Combur dipstick test (Roche, Basel, Switzerland). Female and male urine was pooled separately and filtered sterile. For experiments, the female and male urine pools were mixed in a 1:1 ratio, here designed as pooled human urine.

For infections of urothelial microtissues, kde4498 was inoculated from an LB agar plate into LB medium and incubated overnight at 37°C. The culture was then diluted 1:50 in fresh LB medium and incubated for an additional 48 hours at 37°C. Antibiotic selection (30 μg/mL gentamicin) was applied throughout these steps to maintain plasmid expression. On the day of infection, the stationary-phase culture was diluted in pooled human urine to an optical density OD_600_ of 0.2, without antibiotic selection, and incubated with agitation at 150 rpm for 2 hours at 37°C until the culture reached the exponential phase. The bacteria cultures were then further diluted in urine to 10^6^ CFU mL^-1^.

The urine in the lower chamber of the microtissue-containing transwell (Fig. 4f) was replaced with 0.4 mL of the bacterial-urine infection mixture. Samples were incubated for 4 h at 37°C and 5% CO_2_. The transfected urothelial microtissue transwells were then transferred from the standard 24-well plate to a custom-made imaging plate, thereby replacing the infected urine with fresh urine. The custom-made plate for high-resolution imaging of the inserts was fabricated using a Prusa i3 MK3S+ 3D printer (PRUSA Research, Prague) with GreenTEC Pro filament (Extrudr, Lauterach). The printed well-plate body was bonded to ThermalSeal RTS™ sealing film (Z734438, Sigma-Aldrich), which served as a permeable substrate.

Imaging was performed using a Nikon Spinning disk CSU-W1 microscope equipped with a 40x water-immersion objective (numerical aperture 1.15) at 37°C and 5% CO_2_ in three imaging channels: Cy5 (for bladder tissue cells), Cy3 (for mScarlet), GFP (for FAST:HMBR). HMBR was added to the lower chamber at a final concentration of 5 μM and incubated for 1 h on the microscope. Three positions per transwell were imaged +/-8 μm from the selected plane with step of 0.3 μm. Trimethoprim was then added to a final concentration of 100 μM for 10 min, after which imaging was repeated for the same positions with the same settings. The addition of trimethoprim into the urine of the microtissue model can cause minor shifts in the *xyz*-position of the imaging plate, which can introduce xyz-shifts in the imaging field.

### Image analysis of bladder tissue model infections with UPEC

To quantify biosensor signal, we prepared training data for instance segmentation of individual cells based on the mScarlet signal. We used StarDist^44^ and StarDist OPP^45^ as segmentation models. The input data was normalized based on the 1 %- and 99.8 %-percentiles. Training was performed on the sciCORE cluster with a batch size of (32, 128, 128) and 64 rays. For each position, the two timepoints were segmented independently. After segmentation, only objects with an aspect ratio above 2 were used as correct cell segmentations and further processed. For the FAST:HMBR and the mScarlet channel, the background values were determined as median of all non-segmented pixels for each position and timepoint individually. For each cell, the median FAST:HMBR and mScarlet within the respective segmentation mask was calculated and the ratio (FAST:HMBR_i_ - FAST:HMBR_bg_)/(mScarlet_i_ - mScarlet_bg_) was calculated for every cell with intensities FAST:HMBR_i_ and mScarlet_i_. FAST:HMBR_bg_ and mScarlet_bg_ are the respective background medians.

## Ethics declaration

The use of pooled, sterile-filtered human urine from anonymized healthy volunteers was approved by the relevant ethics committee (Ethikkommission Nordwest-und Zentralschweiz, Project-ID 2023-00592).

## Competing interests

The authors declare no competing financial interest, and no non-financial competing interest.

**Supplementary information** is available in the online version of the paper.

**Correspondence and requests for materials** should be addressed to Urs Jenal and Knut Drescher.

## Acknowledgements

We thank Stella Stefanova and Svitlana Malysheva from the FACS core facility of the Biozentrum (University of Basel) for assistance, teaching, and troubleshooting of FACS-related matters, Konstantin Neuhaus for help with microscopy, Daniel K. H. Rode for help with biofilm assays and BiofilmQ image analysis. Fabienne Hamburger, Kerstin Strenger and Dibya Saha for help with plasmid cloning and strain construction, Raphael Dias Teixeira for advice on preparation of structural models, and Nichole Wespe for support in data management. We are also grateful to Pablo R. Fuentes and Hannes Link for advice and feedback on the project, and to Jörg Vogel and Dirk Bumann for providing *Salmonella enterica* strains. This project was supported by the National Center of Competence in Research (NCCR) AntiResist funded by the Swiss National Science Foundation (51NF40_180541 to U.J. and K.D.), the Swiss National Science Foundation project grant (310030_208107 to U.J.), and by the Swiss National Science Foundation Consolidator Grant (TMCG-3_213801 to K.D.). Additionally, this work was supported by the Deutsche Forschungsgemeinschaft via the Priority Programme SPP2389 (DR 982/6-1 to K.D.), European Union’s Horizon 2020 research and innovation program through the Marie Skłodowska-Curie grant agreement No. 955910 (PHYMOT consortium), an EMBO Postdoctoral Fellowship (ALTF 563-2023 to G.R.), and an International Human Frontier Science Program Organization (HFSPO LT0017/2023-L to C.F.).

## Author contributions

D.F., A.K., K.D. and U.J. designed the project, interpreted the data, and co-wrote the paper with help of all authors. D.F. and A.K. developed and optimized antibiotic biosensors and generated bacterial strains. A.K. performed *in vitro* characterization of biosensors and analyzed the resulting data. N.B.-N. performed image data analysis and statistical calculations. D.F. and G.R. performed and analyzed *S. enterica* infection of macrophages experiments. R.P.J. and T.M. purified the proteins and contributed materials. I.S., A.K., C.F., A.H. and C.D. established and performed UPEC infections of the bladder tissue model. E. J.-S. and D.F. performed BiofilmQ analysis. E.M. synthesized HMBR and HBR-3,5-DOM and contributed materials. S.K. and S.T.S. organized the sampling of human urine. U.J. and K.D. supervised and coordinated the project.

## Supplementary Information

**Suppl. Fig. 1.**
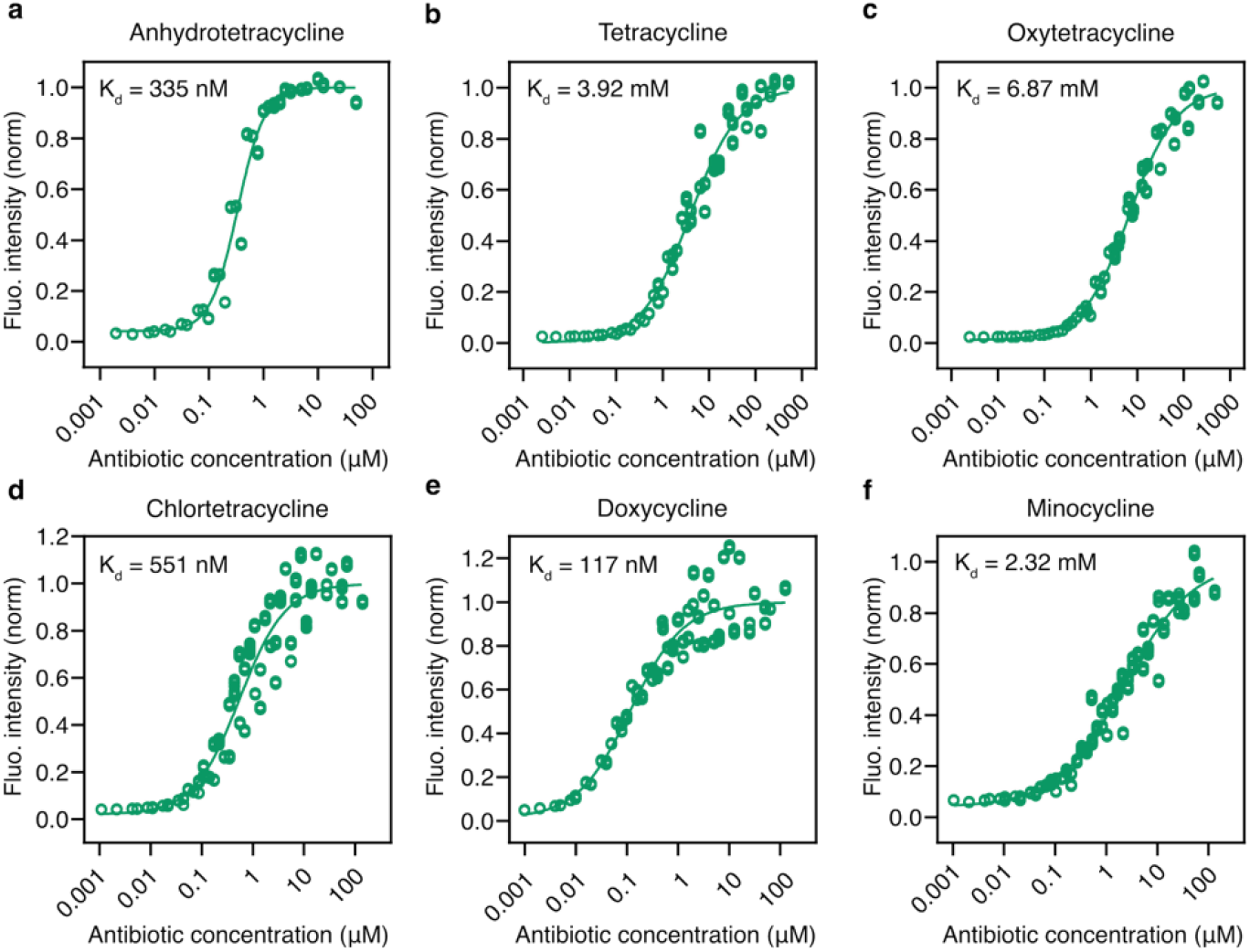
Dose-response curves of TetFABS biosensor with different antibiotics of the class of tetracyclines. **a-f**, The different affinities (K_d_) are shown for each dose-response curve. The K_d_ was obtained by fitting dose-response curves. Data were normalized to the maximal fluorescent intensities determined by fitting (see Methods). Note that the dose-response curve in panel a was prepared from some of the same data shown in Fig. 1k for the 497 nm curve. Data points represent pooled data from n=3 (panel a), n=5 (panels b and c), n=6 (panels d-f) independent dilution series with six (panel a) or eight (panels b-f) repeated measurements for each concentration at steady state.

**Suppl. Fig. 2.**
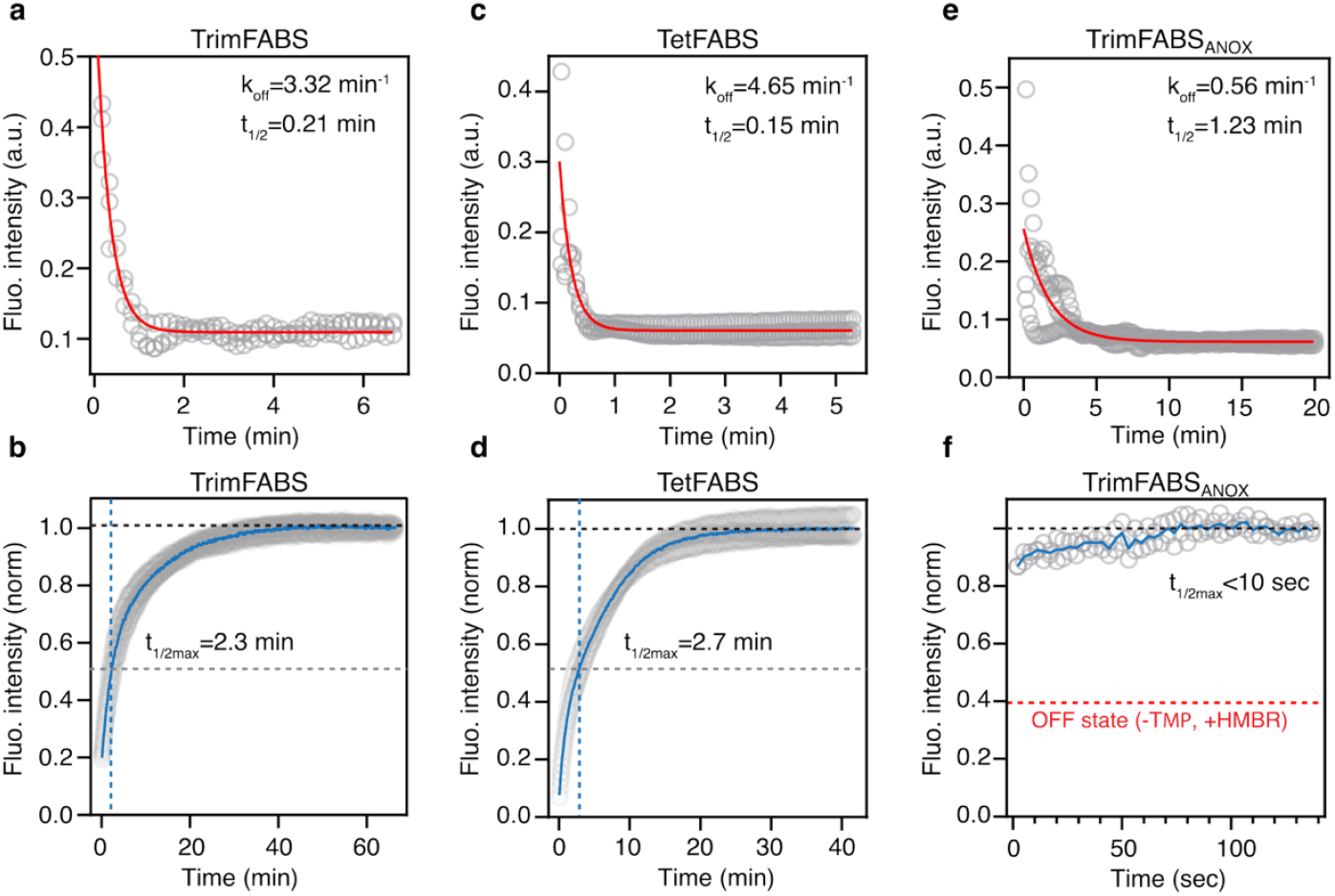
On/off kinetics of all biosensors developed in this study. Panels **a**,**c**,**e** show the off-kinetics, listing the dissociation rate constant k_off_ and the half-life t_1/2_ for each sensor. Panels **b**,**d**,**f** show the on-kinetics, listing the time until half of the maximum fluorescence has been achieved (t_1/2max_) for each sensor. **a**, Dissociation kinetics of the TrimFABS sensor and trimethoprim in the presence of 250 μM NADPH, as determined by the “dissociation by dilution” method. **b**, Association kinetics of TrimFABS sensor in the presence of 250 μM NADPH upon addition of saturating concentrations of trimethoprim (10 μM). **c**, Dissociation kinetics of theTetFABS sensor and doxycycline as determined by the “dissociation by dilution” method. **d**, Association kinetics of sensor TetFABS upon addition of saturating concentrations of doxycycline. **e**, Dissociation kinetics of the TrimFABS_ANOX_:HMBR sensor and trimethoprim in the presence of 5 μM HMBR as determined by the “dissociation by dilution” method. **f**, Association kinetics of the TrimFABS_ANOX_ sensor in the presence of 5 μM HMBR upon addition of saturating concentrations of trimethoprim (10 μM). Individual data points are shown as grey circles. The blue lines in panels b, d and f represent the mean of independent replicates. For dissociation kinetics, the dissociation rate constant k_off_ and the half-life t_1/2_ were derived from a fit (red lines) according to the “Dissociation - One phase exponential decay” model (see Methods). In panel f, gray points show data from n=2 independent replicates. In panels a, b, c, d, e the gray data points correspond to n=3 independent replicates.

**Suppl. Fig. 3.**
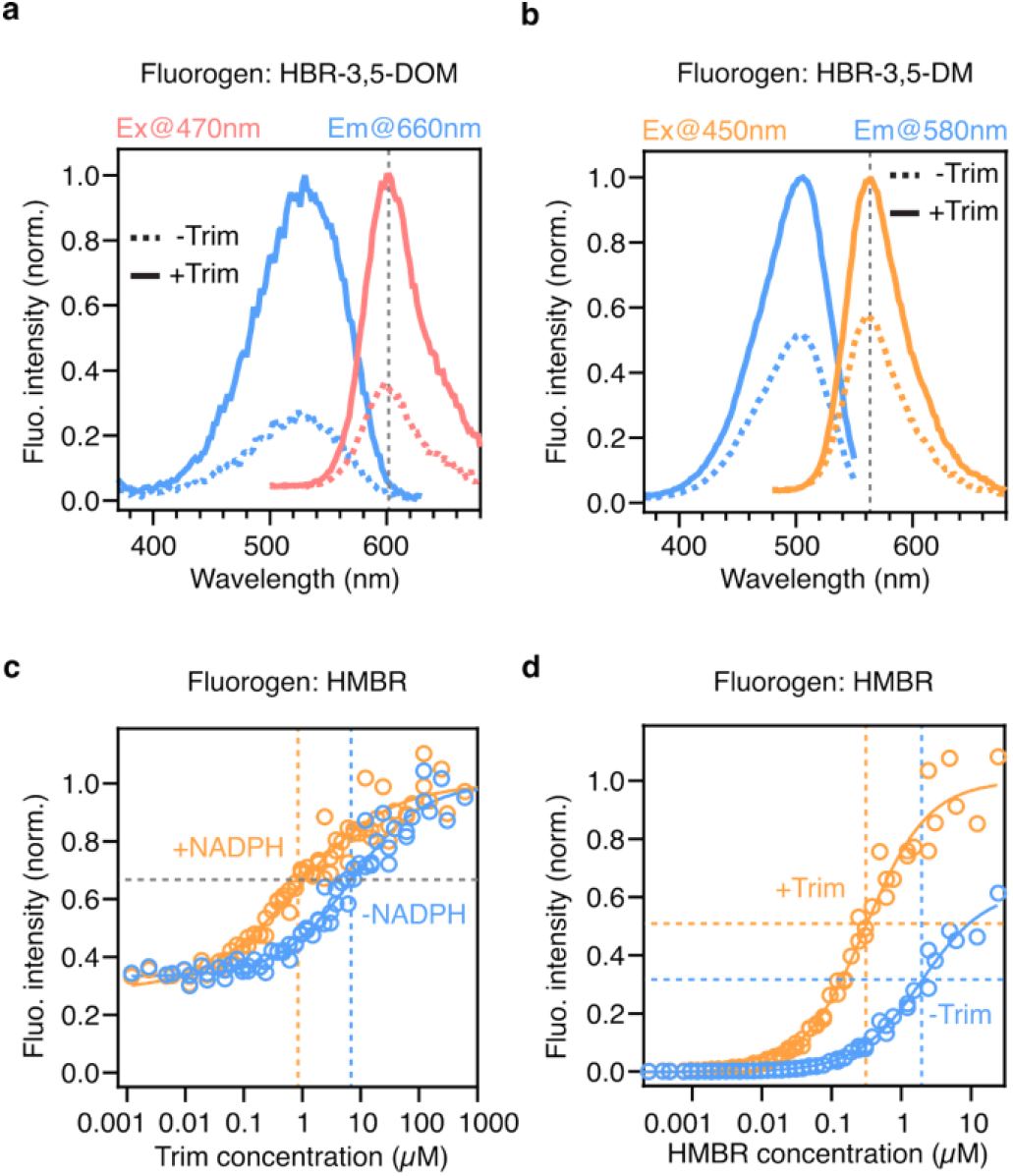
TrimFABS_ANOX_ *in vitro* charachterization. **a**, Excitation and emission spectra of TrimFABS_ANOX_ with (solid lines) or without (dashed lines) Trim, always in the presence of the fluorogen HBR-3,5DOM. Red lines are the emission spectra that result from excitation at 470 nm. Blue lines are the excitation spectra that result from emission measurements at 660 nm. **b**, Excitation and emission spectra of TrimFABS_ANOX_ with (solid lines) or without (dashed lines) Trim and the fluorogen HBR-3,5DM. **c**, Trim dose-response curves of the TrimFABS_ANOX_ sensor with (250 μM) or without NADPH (excitation at 488 nm and emission at 545 nm). Data points represent pooled data from n=6 independent dilution series with six repeated measurements for each concentration at steady state. Dotted lines indicate ligand concentrations at half-maximal response, i.e. dissociation constants (K_d_). **d**, HMBR dose-response curves of the TrimFABS_ANOX_ sensor with (250 μM) or without Trim (excitation at 488 nm and emission at 545 nm). Data points represent pooled data from n=6 independent dilution series with six repeated measurements for each concentration at steady state. Dotted lines indicate ligand concentrations at half-maximal response, i.e. dissociation constants (K_d_).

